# Molecular Insights into Cholesterol Concentration Effects on Planar and Curved Lipid Bilayers for Liposomal Drug Delivery

**DOI:** 10.1101/2025.04.06.647491

**Authors:** Ehsaneh Khodadadi, Ehsan Khodadadi, Parth Chaturvedi, Mahmoud Moradi

## Abstract

Liposomal carriers provide a flexible and effective strategy for delivering therapeutics across a broad spectrum of diseases. Cholesterol is frequently included in these systems to improve membrane rigidity and limit permeability. Despite its widespread use, the optimal cholesterol-to-lipid proportion for achieving stable and efficient liposome performance remains to be fully determined. In this work, we apply all-atom molecular dynamics simulations to explore how different cholesterol concentrations influence the structural and dynamic characteristics of 1,2-distearoyl-sn-glycero-3-phosphocholine (DSPC) bilayers, considering both planar and curved membrane geometries. Bilayers with cholesterol molar ratios of 0%, 10%, 20%, 30%, 40%, and 50% were simulated, and key biophysical parameters including area per lipid (APL), membrane thickness, leaflet interdigitation, and deuterium order parameters (SCD) were analyzed. In planar bilayers, increasing cholesterol concentration led to a progressive decrease in APL from approximately 60 ^°^A^2^ to 40 ^°^A^2^, accompanied by increased membrane thickness and lipid ordering, consistent with cholesterol’s classical condensing effect. In contrast, curved bilayers exhibited a cholesterol-induced expansion effect, particularly in the inner leaflet, where APL increased from approximately 60 ^°^A^2^ to 90 ^°^A^2^ with rising cholesterol levels. SCD profiles showed that cholesterol enhanced tail ordering up to 40% concentration, beyond which the effect plateaued or slightly declined, suggesting structural saturation or packing frustration. Membrane thickness displayed a monotonic increase in planar bilayers but followed a nonlinear trend in curved systems due to curvature-induced stress. These findings highlight that cholesterol’s influence on membrane properties is highly dependent on bilayer geometry and asymmetry. While planar bilayers exhibit predictable responses, curved systems reveal nonclassical behaviors that challenge traditional models of cholesterol-lipid interactions. This work provides molecular-level insights and establishes a computational framework for the rational design of liposomal systems, emphasizing the need to account for curvature and asymmetry in membrane engineering.

## Introduction

Liposomal drug delivery systems have gained significant attention due to their ability to enhance drug stability, reduce toxicity, and provide controlled release.^1^ The physicochemical behavior of liposomes is strongly governed by their lipid composition, with cholesterol playing a central role in modulating key membrane characteristics. It significantly contributes to the regulation of bilayer integrity, permeability, and mechanical resilience.^2,3^ As a fundamental constituent of biological membranes, cholesterol affects lipid arrangement, enhances membrane fluidity, and influences the functionality of associated membrane proteins. ^4,5^ It interacts with phospholipids to modulate bilayer structure, promoting phase separation and the formation of ordered lipid domains.^6^ Cholesterol acts as a condensing agent, reducing the lateral area per lipid while increasing bilayer thickness, lipid tail ordering, and overall mechanical strength.^7,8^ This effect leads to decreased permeability, which is critical for maintaining drug encapsulation and controlled release in liposomal formulations.

At different concentrations, cholesterol can either stabilize or disrupt lipid bilayers, influencing their structural organization and dynamic properties.^9,10^ Additionally, cholesterol’s interaction with phospholipids such as DSPC determines key biophysical properties, including leaflet interdigitation and lipid order.^11,12^ The degree of lipid interdigitation, which refers to the overlap of lipid tails between opposing bilayer leaflets, is another important factor influenced by cholesterol concentration.^13^ While cholesterol enhances membrane stability, excessive amounts may alter lipid organization and impact the bilayer’s ability to function as a drug carrier.^9,14^ Furthermore, liposome stability is affected by several factors, including lipid composition, storage conditions, and interactions with external molecules.^15,16^ In drug delivery applications, maintaining long-term stability is critical, as unstable liposomes may lead to premature drug leakage and reduced therapeutic efficacy.^16–18^ Achieving an optimal cholesterol-to-lipid ratio is essential to balance membrane rigidity and permeability, ensuring effective drug encapsulation and release.

DSPC is widely incorporated into liposomal drug delivery systems due to its high gel-to-liquid crystalline phase transition temperature, which contributes to the formation of robust and stable bilayer structures.^19,20^ The presence of cholesterol within DSPC bilayers plays a critical role in modulating their structural and dynamic properties, including lipid chain packing, bilayer thickness, and overall membrane ordering—all of which are essential factors influencing encapsulation capacity and controlled drug release.^21^ While planar bilayers provide insights into basic membrane behavior, bent bilayers are more representative of liposomal curvature, making their study essential for a complete understanding of cholesterol’s role in liposome design.^22^ Due to the challenges of resolving lipid bilayer structures experimentally, molecular dynamics (MD) simulations serve as a powerful tool for investigating the molecular interactions that govern membrane properties.^23^ MD simulations allow for the precise characterization of lipid-cholesterol interactions at the atomic level, complementing experimental studies and reducing the cost of material-based liposome synthesis.

This work employs all-atom molecular dynamics simulations to explore how varying cholesterol concentrations influence the structural properties of DSPC bilayers. We simulate cholesterol molar ratios of 100%, 80-20%, 70-30%, 60-40% and 50-50% in both planar and bent bilayers, quantifying key properties such as lipid packing, bilayer thickness, and segmental order parameters. By comparing these systems, we aim to determine the optimal cholesterol-lipid arrangement for achieving stable and efficient liposomal formulations. These simulations provide molecular-level insights into the structural and dynamic behavior of DSPC-cholesterol bilayers, establishing a computational framework for the rational design of liposomal drug delivery systems.

## Methods

### Simulation Systems

To investigate the influence of cholesterol concentration and membrane geometry, we constructed planar and curved bilayer systems consisting of DSPC mixed with cholesterol at molar ratios of 100%, 80:20, 70:30, 60:40, and 50:50. All systems were built using the CHARMM-GUI Membrane Builder.^24^ Each system was fully solvated with TIP3P water molecules and neutralized with 0.15 M NaCl to replicate physiological conditions. The simulation boxes measured approximately 82×82×85 ^°^A^3^ for planar bilayers and 127×127×104 ^°^A^3^ for curved bilayers. Energy minimization was performed for 10,000 steps using the conjugate gradient algorithm, followed by equilibration consisting of a 1 ns NVT phase and a 5 ns NPT phase, both under periodic boundary conditions.

All-atom MD simulations were conducted using NAMD 2.14^25^ in combination with the CHARMM36 additive force field,^26^ which is tailored for phospholipid and sterol systems. Temperature was regulated at 310 K using a Langevin thermostat with a damping coefficient of 0.5 ps*^−^*^1^. Pressure control was achieved at 1 atm through the Nośe–Hoover Langevin piston scheme. Non-bonded interactions employed a smoothed cutoff between 10 and 12 ^°^A, and long-range electrostatic interactions were computed via the particle mesh Ewald (PME) algorithm.^27^ Each simulation was extended for a total duration of 200 ns.

To generate curved bilayer systems, we employed a semi-automated workflow that integrated molecular structure preparation in VMD with a curvature induction protocol. Systems were centered at the origin and stripped of water and ions. A new solvation box with 0.15 M NaCl was added, and lipid tail atoms were selected to define the region where curvature-inducing potentials using Grid Forces feature of NAMD^28^ would later be applied. Bilayer thickness was computed from the center-of-mass difference between the upper and lower leaflets along the membrane normal. Curvature was imposed iteratively by adjusting an external potential field; at each step, the bending radius was recalculated based on system geometry, and updated simulation input files were generated. This procedure was repeated until the desired curvature radius of 175 ^°^A was reached. All steps were automated in a dedicated directory to ensure consistency across systems.

Structural analysis and visualization were carried out using VMD,^29^ employing its builtin plugins to evaluate membrane characteristics, including thickness, interdigitation, and lipid tail ordering (*S_CD_*).^30^ Interdigitation between lipid acyl chains of opposing leaflets was quantified using the Interdigitation Tool from the MEMBPLUGIN extension in VMD. This tool calculates the extent of leaflet interpenetration based on three descriptors: the mass overlap fraction (*I_ρ_*), which quantifies the correlation between the mass density profiles of the upper and lower leaflets (ranging from 0 for no overlap to 1 for complete overlap); the width of the overlap region (*w_ρ_*), which represents the thickness of the bilayer region where significant mass overlap occurs; and the contact fraction (*I_c_*), defined as the proportion of atoms from one leaflet that are within 4 ^°^A of atoms in the opposite leaflet. All interdigitation metrics were averaged over time across the full equilibrated trajectory using default parameters and automatic bilayer center detection.

Membrane thickness was computed using the Membrane Thickness Tool from MEMBPLUGIN, which calculates the distance between phosphorus atoms of opposing DSPC headgroups projected onto the *xy*-plane. The average bilayer thickness was recorded over time, and thickness maps were generated to visualize spatial variations and potential domain formation across the membrane. This analysis was applied to both pure DSPC bilayers and DSPC:cholesterol mixtures.

The area per lipid (APL) for planar bilayers was calculated using the MEMBPLUGIN Area per Lipid Tool. This plugin uses Voronoi tessellation to estimate the area surrounding each lipid, based on the positions of headgroup atoms projected onto the membrane plane. MEMBPLUGIN automatically selected three atoms near the glycerol backbone of DSPC to define each lipid’s area contribution. The tool generated both time-resolved APL values and histograms to assess lipid packing as a function of cholesterol concentration.

For curved bilayers, the APL was calculated from MD trajectories using the MDAnalysis Python library. Lipid headgroups were identified using the residue name DSPC and atom name P. A central cubic region (100 ^°^A × 100 ^°^A × 100 ^°^A) was selected, and headgroups were classified into outer or inner leaflets based on their *z*-coordinates. The surface area of this patch was calculated by projecting the cube onto a spherical surface using:

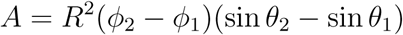

where *R* is the leaflet-adjusted radius (outer or inner), and *ϕ*, *θ* represent angular boundaries. APL was obtained by dividing the projected area by the number of lipid headgroups in each leaflet, frame-by-frame. The results were saved in CSV format and visualized to compare time-dependent behavior.

The bilayer thickness for curved systems was computed using a custom MDAnalysis script. Lipid headgroup positions were projected onto the *xy*-plane and grouped into 50 concentric annular bins extending up to 100 ^°^A from the vesicle center. Radial distances from the spherical center were calculated, and the most prominent peaks—corresponding to inner and outer leaflets—were identified using the find peaks function from scipy.signal, with a minimum peak prominence of 2. The difference between peak positions defined the local thickness per bin. These values were averaged across all bins and timepoints to produce a complete thickness profile over the trajectory.

The deuterium order parameter (*S_CD_*) was calculated using the Membrane SCD Tool in MEMBPLUGIN. This parameter quantifies the orientational ordering of lipid tails relative to the membrane normal (*z*-axis), providing insight into bilayer packing and phase behavior. For each methylene group along the two acyl chains (sn-1 and sn-2) of DSPC, *S_CD_* was calculated using the following equation:

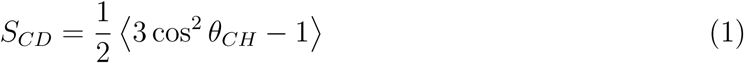

where *θ_CH_* is the angle between the membrane normal and the C–H bond vector, and the brackets denote averaging over time and ensemble. Default parameters were used, and analysis began after equilibration. The output included *S_CD_* profiles along each chain, allowing comparison between cholesterol-containing and cholesterol-free systems.

## Results and Discussion

### Area Per Lipid Analysis and the Role of Cholesterol and Curvature

The area per lipid analysis shows that both cholesterol concentration and membrane curvature play a key role in determining lipid packing. In planar bilayers, cholesterol leads to a condensing effect, where lipids become more tightly packed. In contrast, curved bilayers display the opposite trend, where cholesterol contributes to membrane expansion, particularly in the inner leaflet. These patterns are consistently observed across both symmetric and asymmetric systems in time-series data and replica-averaged results (Figure 1 and S1,S2).

**Fig. 1.**
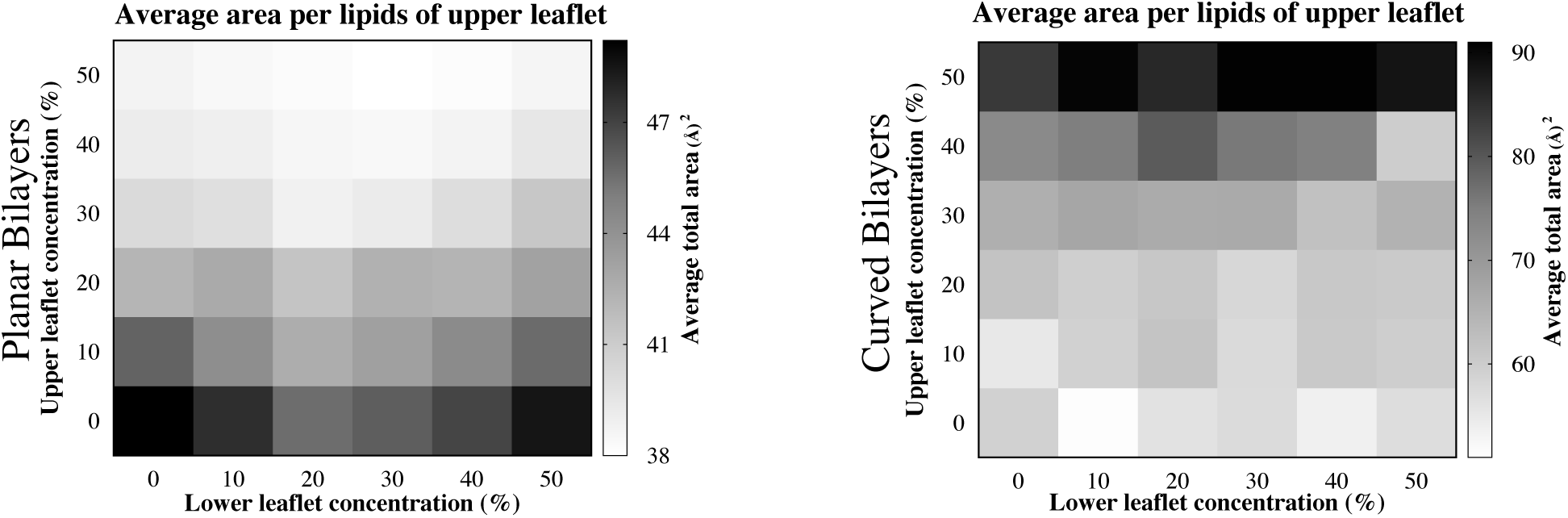
Lipid area and cholesterol concentration effects on membrane structure. Time evolution of the lipid area for the upper and lower leaflets over a 200 ns simulation, illustrating the impact of varying cholesterol distributions. Each line corresponds to a specific cholesterol condition, labeled as X–Y% Chol, where the first number represents the cholesterol concentration in the upper leaflet, and the second number represents the cholesterol concentration in the lower leaflet. Heatmap of the average lipid area in the upper leaflet as a function of cholesterol concentration in both leaflets. Darker regions indicate lower lipid area, while lighter regions correspond to higher values, demonstrating how cholesterol asymmetry influences membrane packing.

In planar bilayers with equal cholesterol in both leaflets, increasing cholesterol from 0% to 50% results in a steady decrease in area per lipid, from approximately 60 ^°^A^2^ to around 40 ^°^A^2^. This effect is seen equally in both leaflets since planar membranes lack curvature-induced asymmetry. During simulations, the area per lipid drops rapidly within the first 25 nanoseconds, reflecting quick membrane relaxation and cholesterol-driven ordering, before leveling off. Heatmaps support these findings, showing progressively darker regions with increasing cholesterol. These trends are in agreement with previous studies that describe how cholesterol inserts into disordered bilayers, promotes lipid tail ordering, and fills the space left by surrounding lipids. These observations are in line with previous atomistic simulations, which demonstrated that cholesterol induces a strong condensing effect in DOPC bilayers by compressing lipid acyl chains in the sterol ring region.^31^

In asymmetric planar bilayers, where the upper leaflet contains no cholesterol and the lower contains increasing concentrations, the area per lipid in the upper leaflet first decreases up to 20% cholesterol. This suggests tighter lipid packing caused by interleaflet coupling. Beyond 20% cholesterol, the area per lipid in the upper leaflet begins to rise again, likely due to mechanical stress or local buckling as leaflet asymmetry increases. Meanwhile, the lower leaflet shows a consistent decrease in area per lipid from approximately 55 ^°^A^2^ at 10% cholesterol to about 40 ^°^A^2^ at 50%. This behavior confirms the direct condensing effect of cholesterol on the leaflet it occupies. A similar trend is observed when cholesterol is fixed in the upper leaflet and varied in the lower, highlighting the influence of leaflet asymmetry and mechanical coupling. In planar systems, configurations such as 10% cholesterol in the upper leaflet and 0% in the lower are nearly identical to the reverse configuration. This symmetry reflects the balanced mechanical environment of flat membranes. This is consistent with previous findings showing that cholesterol exerts a stronger condensing effect in polyunsaturated bilayers and that area per lipid does not follow simple additive trends due to complex lipid–cholesterol interactions and domain formation at high cholesterol concentrations.^32^ These structural trends are consistent with coarse-grained simulations comparing DSPC and DPSM liposomes, which showed that DSPC lipids form more stable, spherical bilayers due to their cylindrical shape and smaller headgroup, contributing to tighter lipid packing and reduced curvature strain.^33^

In curved bilayers, cholesterol produces a very different outcome (Figure 1 and S1). As the cholesterol concentration increases, whether symmetrically or asymmetrically, the area per lipid also increases, especially in the inner leaflet. In the symmetric 50:50 cholesterol system, the average area reaches about 90 ^°^A^2^ compared to 60 ^°^A^2^ in the cholesterol-free system. This expansion is driven by curvature-induced stress. The outer leaflet stretches while the inner leaflet compresses, and this mechanical effect overrides the usual condensing behavior of cholesterol. Heatmaps clearly show this contrast, with curved systems displaying a near-linear increase in area per lipid and greater differences between leaflets, especially at higher cholesterol levels.

In asymmetric curved bilayers, where cholesterol is added only to the inner leaflet, the area per lipid in the outer leaflet increases with cholesterol up to 30%, then decreases slightly at 40%, and increases again at 50%. This irregular pattern likely reflects the redistribution of stress caused by curvature. At the same time, the area per lipid in the inner leaflet increases steadily from the lowest value at 10% to about 90 ^°^A^2^ at 50%. This is in sharp contrast to planar systems, where cholesterol always reduces the area per lipid. The expansion observed in curved systems is likely caused by geometric constraints. The inner leaflet has a smaller surface area and experiences radial compression, so cholesterol may trigger lipid rearrangement that paradoxically leads to increased spacing.

Similar patterns emerge when cholesterol is added to the outer leaflet and gradually introduced to the inner one. Across all curved systems, heatmaps and averaged results confirm these trends and show a pronounced asymmetry between the leaflets. Curvature appears to magnify the impact of leaflet composition and cholesterol distribution. One key insight is that curvature introduces directional asymmetry. In curved systems, placing 10% cholesterol in the outer membrane layer and none in the opposite one is not equivalent to reversing this distribution. This difference arises from the distinct mechanical environments of the inner and outer leaflets. The outer leaflet tends to expand more easily, while the inner one is constrained by geometry. This directional dependence helps explain why lipid packing patterns diverge under curvature and is especially important in cholesterol-rich vesicles or nanodiscs. These observations are also consistent with atomistic simulations of plasma membranes under controlled curvature, which showed that membrane bending causes leaflet-specific changes in area per lipid, with the convex leaflet expanding and the concave leaflet compressing, confirming that curvature-induced stress governs lipid packing and overrides classical condensation effects.^34^

### Segmental Order Parameter (SCD) Reveals Cholesterol-Induced Tail Rigidity

Understanding how cholesterol modulates membrane structure is essential for elucidating its role in bilayer stability, domain formation, and lipid packing. One of the most informative indicators of membrane order is the segmental order parameter (SCD), which quantifies the alignment of lipid acyl chains. Higher SCD values reflect increased ordering and rigidity of lipid tails, while lower values indicate enhanced flexibility and disorder. (Figure 2 and S2) present a comprehensive overview of SCD profiles for both sn1 and sn2 chains under various symmetric and asymmetric cholesterol distributions, capturing leaflet-specific behaviors in both planar and curved bilayers. These data provide a systems-level view of how cholesterol concentration and leaflet asymmetry influence bilayer ordering.

**Fig. 2.**
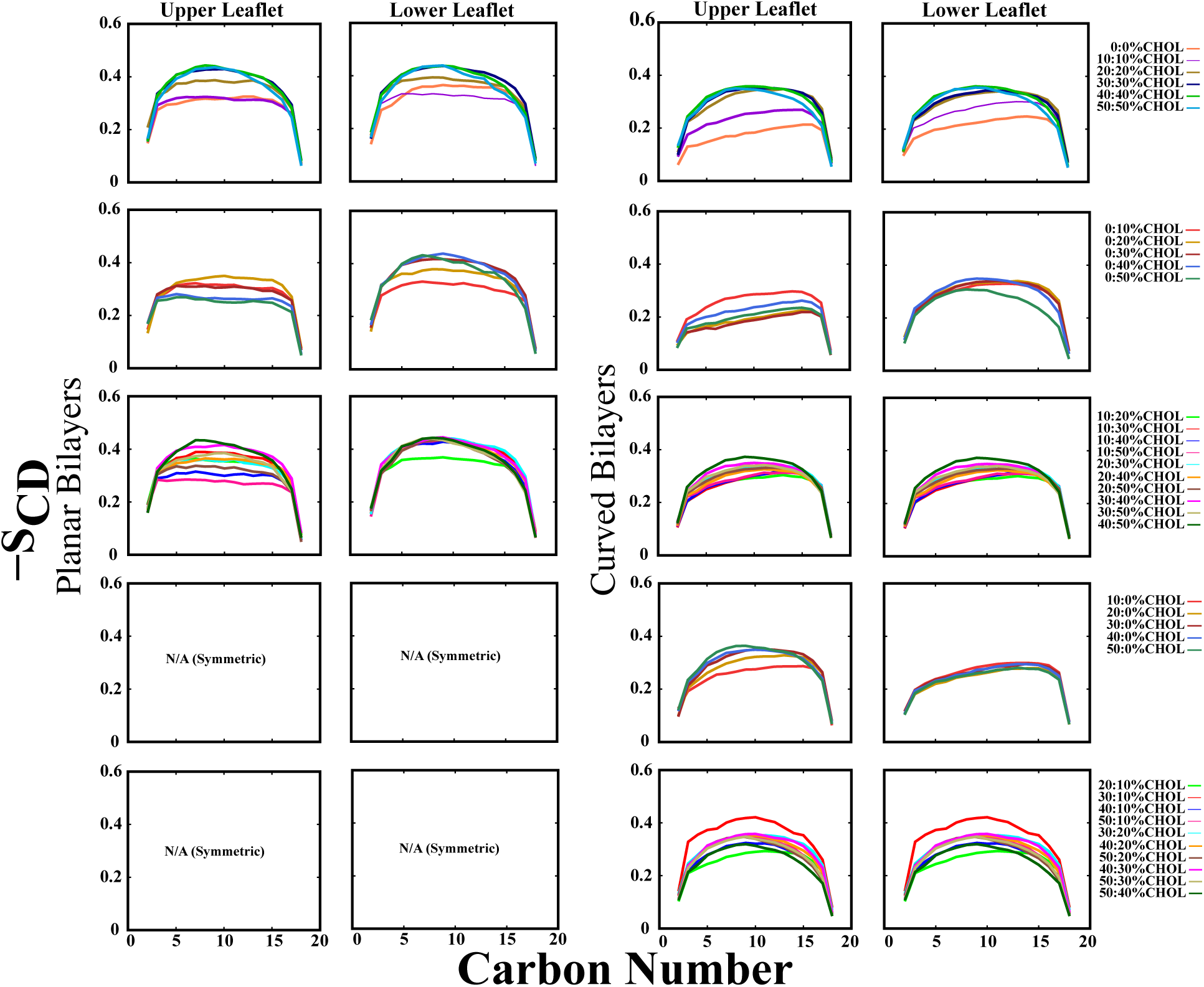
Deuterium order parameters and cholesterol concentration effects on membrane structure. Segmental order parameter (SCD) profiles of the SN2 hydrocarbon chain for the upper and lower leaflets, illustrating the impact of varying cholesterol distributions on membrane ordering. Each line represents a specific cholesterol condition, labeled as X–Y% Chol, where the first number indicates the cholesterol concentration in the upper leaflet, and the second number represents the cholesterol concentration in the lower leaflet. Higher (SCD) values indicate increased lipid tail rigidity, highlighting how cholesterol asymmetry influences membrane order across different bilayer configurations.

At 0% cholesterol, the bilayer is highly flexible and disordered, with SCD values between ∼0.22 and 0.34 (Figure 2 and S2). Increasing cholesterol to 10%–30% leads to a progressive rise in SCD, particularly in the mid-chain region (carbon atoms 5–15), reflecting enhanced packing. Beyond 40% cholesterol, the ordering effect plateaus, and a slight decrease in SCD is observed at terminal carbons, suggesting limited additional ordering and potential structural disruption at higher concentrations. This trend aligns with previous findings where higher concentrations of ordered-phase lipids, such as DSPC, increase acyl chain order due to its gel-phase behavior below the phase transition temperature (*T*_m_ = 328 K). Cholesterol mimics this effect, promoting a transition to a more condensed bilayer. However, at concentrations exceeding 40%, a slight reduction in tail order may signal the onset of packing frustration or phase separation.^35^ The sn2 chain consistently exhibits slightly lower SCD values than sn1, indicating greater flexibility in the second acyl chain, but both chains follow the same trend. Additionally, the upper and lower leaflets exhibit similar ordering patterns in symmetric bilayers, indicating that cholesterol exerts a balanced effect when evenly distributed. To further explore asymmetric cholesterol distribution, we analyzed systems where only the lower leaflet contained cholesterol at 10%, 20%, 30%, 40%, or 50%, while the upper leaflet remained cholesterol-free. In these 0:10% to 0:50% CHOL systems, the upper leaflet remained largely disordered, with SCD values around 0.28–0.32, showing minimal variation and limited evidence of cross-leaflet influence. At 0:20% CHOL, a slight increase in upper leaflet SCD was observed, possibly indicating weak interleaflet coupling, but this effect diminished at higher concentrations. In contrast, the lower leaflet exhibited a strong, cholesterol-dependent increase in order. SCD values rose from ∼0.33 at 0:10% to ∼0.46 at 0:50%, confirming cholesterol’s ability to enhance local lipid packing and reduce tail flexibility. These trends were consistent across both planar and curved bilayers, although curvature slightly reduced the magnitude of SCD due to geometric constraints.

We also examined systems in which cholesterol was present only in the upper leaflet, with the lower leaflet cholesterol-free (10:0%, 20:0%, 30:0%, 40:0%, and 50:0%). Planar bilayers were not analyzed for these cases due to symmetry (e.g., 10:0% is equivalent to 0:10%).However, in curved bilayers, membrane geometry leads to structural differences across the two leaflets. Notably, the outer leaflet, enriched with cholesterol, exhibited a concentration-dependent rise in SCD values, reaching a maximum of approximately 0.42 at 50:0% CHOL. The cholesterol-free lower leaflet remained disordered across all cases (SCD ∼0.25–0.28), reinforcing the localized nature of cholesterol’s effect, even under curvature.

Bilayers with differing cholesterol concentrations in both leaflets, such as 10:50%, 30:40%, or 20:30% CHOL, revealed strong leaflet-specific ordering. The cholesterol-enriched leaflet consistently exhibited higher SCD values (∼0.44–0.46), while the less-enriched leaflet remained less ordered (∼0.30–0.34). In curved membranes, this asymmetry was slightly reduced, suggesting curvature-induced stress redistribution may promote partial interleaflet mechanical coupling. Nonetheless, cholesterol’s effects remained primarily localized to the leaflet in which it resided. These results demonstrate that cholesterol’s impact on lipid tail ordering is highly localized and leaflet-specific. Planar bilayers display more pronounced and symmetric ordering, while curved bilayers show slightly reduced SCD due to geometric frustration. Regardless of geometry, local cholesterol concentration remains the dominant factor governing lipid organization in bilayer membranes.

Interestingly, the plateau and eventual decrease in SCD values at high cholesterol concentrations (e.g., ≥40%) may be attributed to structural frustration within the bilayer. Solidstate NMR studies have shown that while cholesterol induces a significant increase in lipid tail ordering, this effect saturates at intermediate concentrations and can decline beyond a certain threshold. Molugu et al.^36^ demonstrated that cholesterol induces a plateau in the segmental order parameter (SCD) profiles of acyl chains in the liquid-ordered phase, approaching the limit of an all-trans configuration (SCD ≈ −0.5), beyond which further ordering becomes sterically unfavorable. This saturation implies a physical limit to cholesterol’s condensing power, and excessive cholesterol can introduce packing frustration, disrupting uniform lipid alignment and potentially triggering local phase separation. These findings reinforce our observations that SCD values rise with increasing cholesterol but reach a plateau or decline slightly at very high concentrations, consistent with the formation of cholesterol-rich domains that may locally disturb lipid packing.

### Structural Thickening Effects of Cholesterol in Lipid Bilayers

Bilayer thickness was assessed to quantify the structural response of lipid membranes under varying cholesterol concentrations and geometries. Cholesterol’s impact on membrane thickness reveals notable differences between planar and curved bilayers, as shown in time-series plots (Figure X) and replica-averaged heatmaps (Figures S3 and S4). In planar bilayers, increasing cholesterol concentration induces a clear, monotonic thickening effect. The average thickness increases from approximately 48 ^°^A in cholesterol-free systems to over 51 ^°^A at 40% CHOL. A slight decrease at 50% suggests a possible saturation threshold or structural reorganization at high cholesterol content. This trend is consistent with cholesterol’s classical condensing effect, where tighter lipid packing and enhanced acyl chain ordering result in membrane thickening.

In curved bilayers, however, the response is more complex and non-monotonic. Thickness increases up to approximately 31 ^°^A at 40% CHOL, followed by a slight decline at 50%. This behavior likely reflects the interplay between curvature-induced stress, leaflet asymmetry, and cholesterol-driven structural rearrangements. Across all concentrations, curved bilayers remain thinner than planar ones, emphasizing the geometric constraint imposed by curvature. Specifically, the inner leaflet experiences radial compression, while the outer leaflet is stretched, both of which counteract cholesterol’s thickening influence.

In asymmetric planar systems, where the upper leaflet remains cholesterol-free and the lower contains varying CHOL levels, thickness initially increases up to 20–30% CHOL before gradually declining. This non-linear trend suggests that moderate asymmetry enhances ordering and thickening, while excessive asymmetry may induce mechanical imbalance or interleaflet relaxation.

A similarly complex trend is observed in asymmetric curved systems. For instance, with 20% CHOL in the upper leaflet, increasing the lower leaflet CHOL content boosts thickness up to 30%, followed by a reduction at higher concentrations. At 40–50% CHOL in the upper leaflet, curved membranes exhibit multiphasic behaviors, including dips and rebounds, not observed in planar bilayers. These trends are likely governed by radial compression, interleaflet stress redistribution, and localized structural adjustments.

Our observation of non-monotonic thickness behavior is consistent with findings by, ^37^ who identified three distinct cholesterol-induced regimes: a miscible liquid-disordered (Ld) phase at low CHOL content, a domain-registered coexistence of liquid-ordered (Lo) and Ld phases at intermediate levels, and a domain-antiregistered “cholesterolic gel” (Soc) phase at high concentrations. This Soc phase, marked by thread-like aggregates of cholesterol and DPPC, correlates with increased curvature and interleaflet antiregistration, in line with our observed rebound effects and abrupt thickness fluctuations at high CHOL in curved systems. Notably, their MARTINI coarse-grained simulations identified a significant morphological transition near 42% CHOL, matching the inflection point in our curved bilayer results.

This curvature-sensitive behavior is also supported by,^38^ who showed that while cholesterol enhances mechanical strength and bilayer thickness at low-to-moderate concentrations, it leads to steric crowding and weakening at higher levels. Their findings indicate that lipid saturation modulates cholesterol’s structural impact, a result consistent with our observation that symmetric planar membranes exhibit more predictable thickening trends than curved or highly asymmetric systems. Moreover, their lateral density analysis revealed that in unsaturated bilayers, cholesterol tends to localize toward the bilayer center, reducing its effect on interleaflet coupling. This supports our findings of softening in highly curved asymmetric membranes.

The study by^39^ using SAXS experiments further supports our results. They reported that cholesterol does not distribute evenly between leaflets in unilamellar vesicles and that curvature amplifies this asymmetry, resulting in leaflet-specific thickening and reorganization. These effects were more pronounced in thinner, monounsaturated lipid systems, reinforcing our interpretation that curvature and leaflet composition jointly modulate cholesterol’s structural effects. They also highlighted hydrophobic mismatch and curvature stress as key drivers of asymmetry, factors clearly influencing the non-monotonic thickening observed in our simulations.

Additional support comes from,^40^ who showed that cholesterol induces uneven curvature in asymmetric membranes, forming extended flat regions interspersed with sharp bends. Their findings suggest that cholesterol senses local curvature and redistributes accordingly, a phenomenon likely responsible for the multiphasic thickness behavior observed in our curved systems. Although their study included DOPS lipids and ours does not, the mechanistic parallels reinforce the importance of curvature in modulating cholesterol behavior.

Overall, planar bilayers exhibit a relatively linear response to increasing CHOL, with consistent thickening followed by minor relaxation at high concentrations. Curved bilayers, in contrast, show nonlinear, geometry-dependent responses driven by asymmetry, radial stress, and localized structural rearrangements. These results highlight the essential role of membrane curvature in regulating cholesterol’s structural influence, a factor that must be accounted for when designing lipid-based drug delivery systems such as vesicles and nanodiscs.

Our findings are also aligned with,^41^ who used 2D IR spectroscopy to study DLPC vesicles and planar bilayers. They observed cholesterol-induced structural transitions at lower concentrations in vesicles (10–15%) compared to planar bilayers (25–30%), consistent with our early onset of non-monotonic thickness changes in curved systems. Faster structural dynamics in vesicles, attributed to curvature-induced stress and density fluctuations, may explain the observed multiphasic thickness patterns at higher cholesterol levels. They proposed a transition between liquid-disordered and liquid-ordered phases depending on cholesterol content and curvature, supporting our conclusion that structural reorganization occurs in both symmetric and asymmetric curved bilayers. ^41^

These insights reinforce the conclusions of, ^40^ who emphasized that cholesterol’s structural impact is not purely concentration-dependent but closely tied to curvature and spatial redistribution. Altogether, the evidence underscores the complexity of designing cholesterol-rich liposomal systems: membrane geometry, lipid saturation, and cholesterol asymmetry must all be carefully optimized to ensure functional stability and effective drug delivery performance.

### Curvatureand Cholesterol-Dependent Interdigitation in DSPC Bilayers

We analyzed lipid tail interdigitation in symmetric and asymmetric DSPC:CHOL bilayers under both planar and curved geometries. In planar bilayers (left column of Figure 4 and S4), interdigitation remained consistently low and stable across all systems, with values fluctuating around ∼0.09–0.12 ^°^A. Increasing CHOL concentration or introducing leaflet asymmetry did not significantly impact interdigitation in planar systems, indicating that bilayer flatness limits tail interweaving, regardless of CHOL content.

**Fig. 3.**
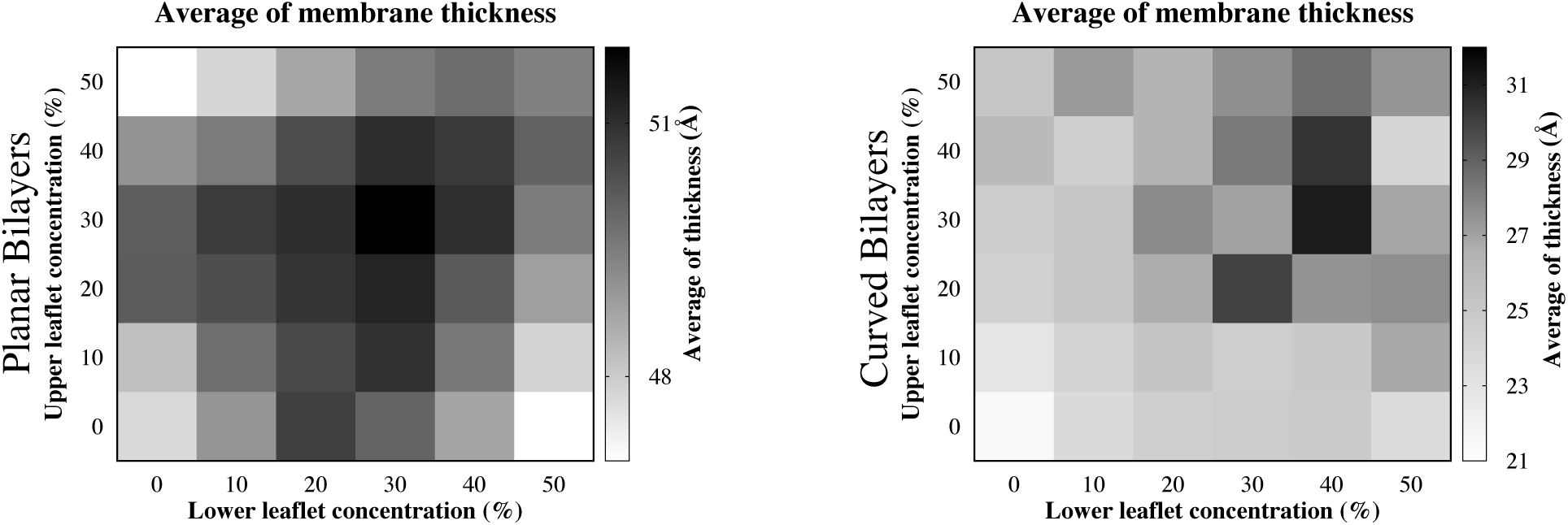
Membrane thickness in planar and curved bilayers. Time evolution and average values for bilayer thickness across various cholesterol distributions. Each condition is labeled as X–Y% CHOL, where the first number denotes the upper leaflet concentration and the second the lower. Heatmaps illustrate how leaflet asymmetry and cholesterol levels affect thickness.

**Fig. 4.**
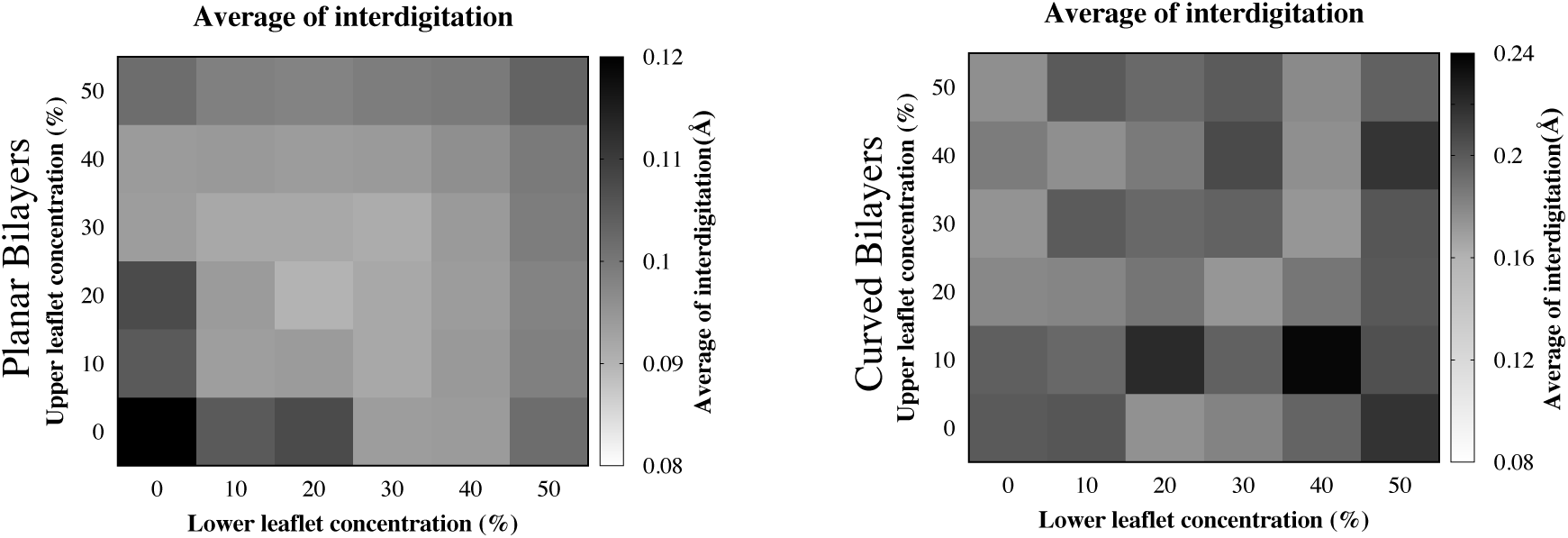
Average lipid tail interdigitation in planar and curved DSPC:CHOL bilayers as a function of upper and lower leaflet cholesterol concentrations. Each heatmap represents the mean interdigitation (in ^°^A) calculated over 200 ns simulations. The left panel shows planar bilayers with relatively low and homogeneous interdigitation values, while the right panel shows curved bilayers with markedly higher and more variable interdigitation, especially at asymmetric and cholesterol-rich conditions. These results highlight the combined influence of membrane curvature and cholesterol asymmetry on interleaflet tail overlap.

In contrast, curved bilayers (right column of Figure 4 and Figure S4) exhibited a clear and progressive increase in interdigitation, particularly in systems with higher CHOL concentrations and asymmetric CHOL distributions. Even symmetric systems such as 40:40 and 50:50 CHOL showed elevated interdigitation, likely due to CHOL-induced condensation enhanced by curvature. The most pronounced effects were seen in asymmetric systems like 0:50, 10:40, and 30:50 CHOL, with final average values reaching up to ∼0.24 ^°^A. These trends suggest that curvature introduces mechanical stress that increases leaflet contact, while asymmetric CHOL distribution leads to lipid tail intrusion from the CHOL-rich leaflet into the opposing side. The result is a synergistic enhancement of interleaflet mixing, driven by both geometry and lipid asymmetry.

Our interdigitation analysis revealed that while planar DSPC:CHOL bilayers maintained consistently low interleaflet mixing, curved bilayers exhibited a progressive and cholesterol-dependent increase in interdigitation, particularly in asymmetric systems. These findings are consistent with detailed molecular dynamics simulations showing that cholesterol promotes significant ordering and condensation of saturated lipid tails like DPPC due to its high planarity and strong van der Waals interactions.^42^ The rigid ring structure of cholesterol aligns well with lipid acyl chains, enhancing membrane order and reducing lipid mobility, especially at higher concentrations. Although those simulations were limited to planar geometries, the observed increase in order and tighter lipid packing directly supports our finding that in curved bilayers, this enhanced ordering may facilitate deeper penetration of lipid tails between leaflets. Additionally, cholesterol’s ordering effect was shown to be greater than that of ergosterol, highlighting its unique role in stabilizing lipid bilayers. ^42^ By extending these insights to curved systems, our results suggest that the combined effects of membrane curvature and asymmetric cholesterol distribution synergistically increase interleaflet contact and promote lipid tail interdigitation, contributing to the structural complexity and functional diversity of cholesterol-rich vesicular membranes.

Our results on interdigitation in planar and curved DSPC:CHOL bilayers align well with findings demonstrating that cholesterol significantly stabilizes DSPC bilayers across a range of propylene glycol (PG) concentrations.^43^ In that study, cholesterol was shown to inhibit the formation of interdigitated lamellar phases that otherwise emerged in DSPC systems exposed to PG, likely by restricting solvent penetration and preserving bilayer thickness. This stabilizing effect was observed up to 60% (w/w) PG, beyond which structural disorder and eventual isotropic phases were noted. Similarly, our data show that in curved geometries, cholesterol’s condensing and ordering effects are enhanced, promoting lipid packing and facilitating interleaflet tail overlap, particularly in asymmetric systems. The prevention of interdigitation in the presence of cholesterol under PG-induced stress supports our interpretation that cholesterol modulates bilayer mechanics by reducing membrane fluidity and enhancing leaflet coupling. These synergistic effects between curvature and cholesterol enrichment contribute to increased interleaflet mixing and bilayer complexity in vesicular systems.

Our observation that curvature enhances lipid tail interdigitation, particularly in asymmetric cholesterol distributions, is further supported by studies of ternary lipid bilayers composed of DOPC, DSPC, and cholesterol on high-curvature substrates.^44^ That work showed that domain coalescence and demixing persisted at higher cholesterol concentrations on curved silica xerogels compared to flat mica surfaces, suggesting that membrane curvature shifts the miscibility threshold and promotes sustained phase separation. The increased interleaflet contact in our curved vesicles likely stems from enhanced lipid segregation and stress-induced rearrangement. It was proposed that curvature either alters cholesterol partitioning or promotes lateral phase demixing by lowering lateral pressure—both of which are consistent with the increased interdigitation we observed.^44^ These insights reinforce the idea that membrane geometry is a crucial regulator of bilayer structural organization and lipid-lipid interactions, especially under conditions of compositional asymmetry.

Our interdigitation findings in DSPC:CHOL bilayers are further supported by experimental results using sum-frequency vibrational spectroscopy (SFVS), which demonstrated that cholesterol alters the physical properties of lipid membranes by introducing packing defects and increasing molecular disorder, even at low concentrations.^45^ This disruption was shown to lower the energy barrier for molecular rearrangements, enhancing bilayer flexibility and dynamic behavior. These observations align with our results showing increased lipid tail interdigitation in curved and cholesterol-rich systems, particularly under asymmetric CHOL distribution. The packing defects and increased fluidity introduced by cholesterol likely facilitate deeper interleaflet contact in curved geometries, where mechanical stress amplifies the disordering effects of CHOL. Together, these findings underscore cholesterol’s dual role as a membrane stabilizer and dynamic modulator, promoting interleaflet mixing in vesicular systems.

These findings contrast with atomistic simulations that reported only ∼2% interdigitation in planar Lo/Ld phase bilayers composed of DOPC, DPPC, and cholesterol.^12^ In that study, interdigitation was not found to be a key contributor to interleaflet coupling and appeared largely independent of CHOL content or phase mismatch, being driven instead by acyl chain properties. Even after altering the area per lipid (APL), interdigitation remained unaffected, suggesting it was not dynamically responsive. In contrast, our curved DSPC:CHOL systems demonstrate that membrane curvature dramatically increases interdigitation, especially under CHOL asymmetry. This reveals a mechanistic difference between planar and curved membranes, where curvature facilitates lipid tail penetration and enables structural behaviors not captured in flat bilayer models. Our results show that interdigitation is minimal in planar DSPC:CHOL bilayers, consistent with prior studies, but becomes significantly enhanced in curved membranes, particularly under asymmetric cholesterol conditions. These findings highlight the importance of considering bilayer geometry and lipid distribution asymmetry when evaluating interleaflet coupling and membrane mechanical properties in systems such as liposomes and curved biological membranes.

## Conclusion

This study reveals that cholesterol’s impact on membrane structure is highly dependent on both its concentration and the geometric configuration of the bilayer. In planar systems, increasing cholesterol consistently induces lipid condensation, characterized by reduced area per lipid, increased membrane thickness, and elevated deuterium order parameters, consistent with its classical role as a condensing agent. These effects are symmetric across leaflets and show clear trends that align with prior experimental and computational studies. In contrast, curved bilayers exhibit markedly different behavior. Cholesterol causes an unexpected expansion in the inner leaflet, increasing the area per lipid and partially opposing the typical condensing effect. This curvature-driven expansion is attributed to radial stress and geometric constraints that dominate over local cholesterol-lipid interactions. The influence of leaflet asymmetry becomes more pronounced under curvature, leading to non-monotonic trends and directional mechanical coupling between leaflets. While cholesterol enhances tail ordering in both geometries, the magnitude and saturation points differ, with curvature slightly reducing the maximum SCD values due to geometric frustration. Membrane thickness trends also diverge between geometries. While planar systems show a monotonic increase with cholesterol content, curved systems exhibit complex, non-linear behaviors influenced by both composition and leaflet-specific stress. These findings highlight the critical interplay between cholesterol concentration, membrane curvature, and compositional asymmetry in governing bilayer structure. Our results demonstrate that classical models of cholesterol condensation do not fully capture its behavior in curved or asymmetric membranes. Curvature imposes directional mechanical stresses that modulate lipid organization and override simple additive effects. These insights are particularly relevant for the rational design of lipid-based nanocarriers, where both curvature and asymmetric composition are inherent. Incorporating these factors is essential for accurately predicting and engineering membrane behavior in biological and therapeutic contexts.

## Supporting Information Available

This article contains Supporting Information.

## Acknowledgement

This research was supported by the National Institute of General Medical Sciences (NIH grant R35GM147423 awarded to M.M.), the National Science Foundation (NSF grant CHE 1945465 awarded to M.M.), and the Arkansas Biosciences Institute. Computational resources were provided by the Texas Advanced Computing Center (TACC) at the University of Texas at Austin (Frontera) through LRAC allocation CHE21003 to M.M. The work also used Stampede at TACC, Expanse at the San Diego Supercomputer Center, and Bridges-2 at the Pittsburgh Supercomputing Center through allocation MCB150129 from the Advanced Cyberinfrastructure Coordination Ecosystem: Services & Support (ACCESS) program, supported by NSF grants #2138259, #2138286, #2138307, #2137603, and #2138296.^46^ Additional computational support came from the Arkansas High-Performance Computing Center, funded by multiple NSF grants and the Arkansas Economic Development Commission.

