## Supporting Information for "Molecular Insights into Cholesterol Concentration Effects on Planar and Curved Lipid Bilayers for Liposomal Drug Delivery"

### List of Figures

S1 Lipid area and cholesterol concentration effects on membrane structure . . . . . . . . . . S3

S2 Mean area per lipid in planar and curved bilayers (upper and lower leaflets) . . . . . . . S4

S3 Segmental order parameter ($SCD$) profiles in planar and curved bilayers . . . . . . . . . . S5

S4 Membrane thickness in planar and curved bilayers . . . . . . . . . . . . . . . . . . . . . S6

S5 Lipid tail interdigitation in planar and curved bilayers . . . . . . . . . . . . . . . . . . . S7

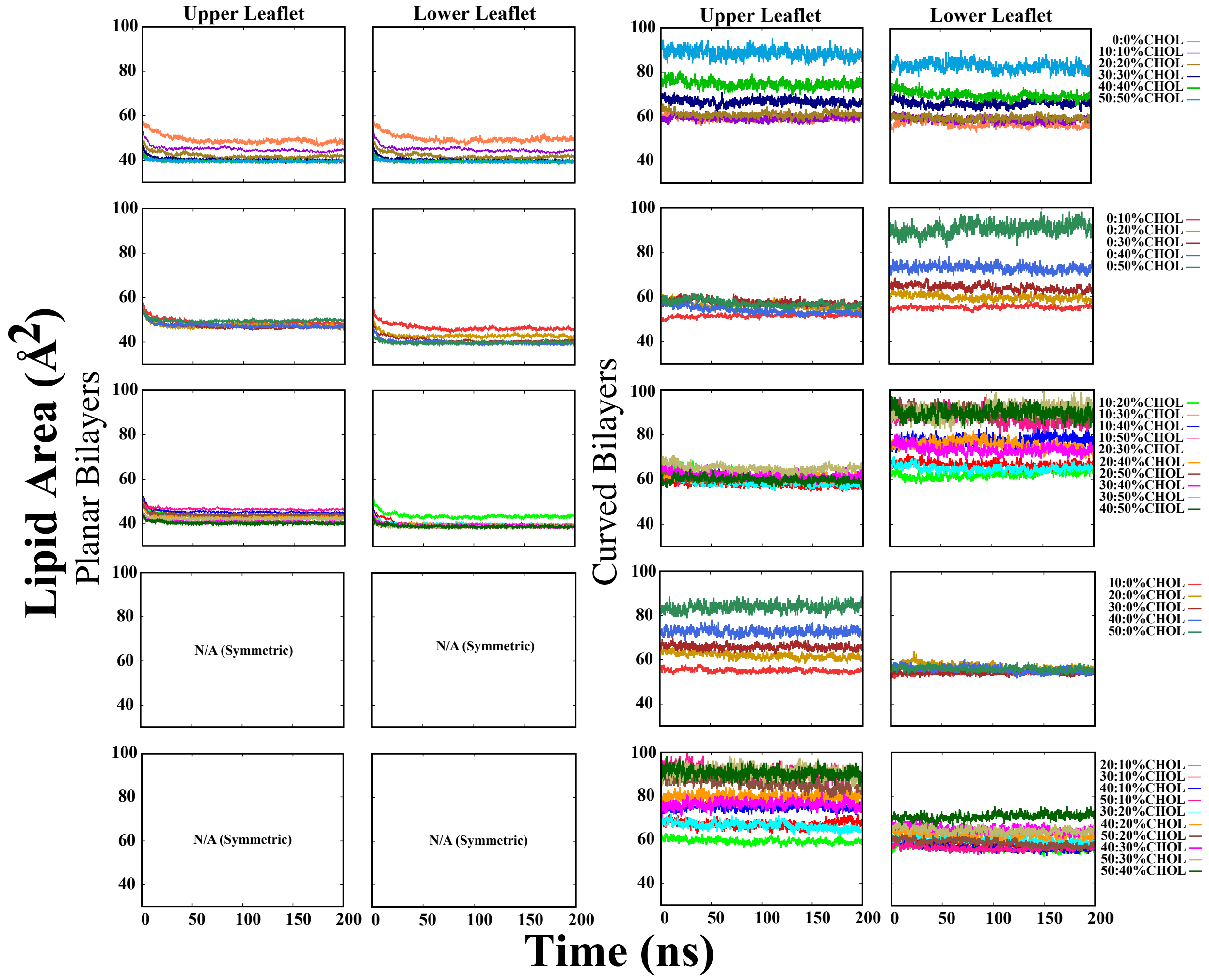


**Fig. S1. Lipid area and cholesterol concentration effects on membrane structure.** Time evolution of the lipid area for the upper and lower leaflets over a 200 ns simulation, illustrating the impact of varying cholesterol distributions. Each line corresponds to a specific cholesterol condition, labeled as X–Y% Chol, where the first number represents the cholesterol concentration in the upper leaflet, and the second number represents the cholesterol concentration in the lower leaflet. A heatmap of the average lipid area in the upper leaflet as a function of cholesterol concentration in both leaflets is also shown. Darker regions indicate lower lipid area, while lighter regions correspond to higher values, demonstrating how cholesterol asymmetry influences membrane packing.

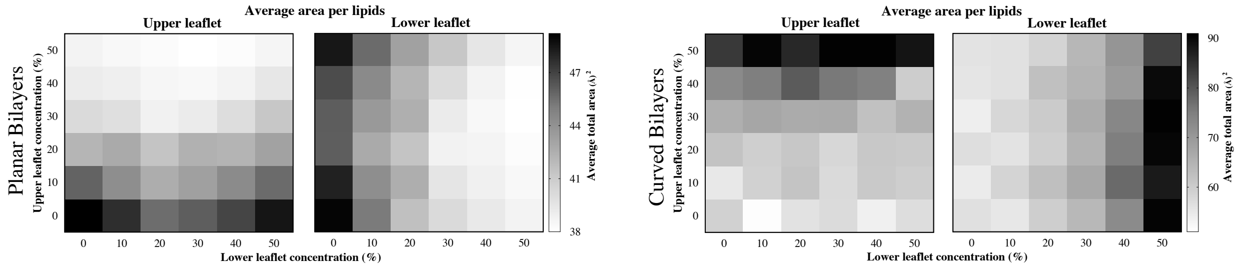


**Fig. S2. Lipid area and cholesterol concentration effects on membrane structure.** Heatmaps showing the lipid area per molecule in the upper and lower leaflets of planar (left) and curved (right) DSPC:CHOL bilayers. Each matrix illustrates how the lipid area varies with cholesterol concentration in the upper (y-axis) and lower (x-axis) leaflets. In planar systems, increasing cholesterol reduces the lipid area, particularly under symmetric conditions, while curved systems exhibit greater sensitivity to cholesterol asymmetry. Darker regions correspond to smaller lipid areas (tighter packing), and lighter regions indicate expanded configurations.

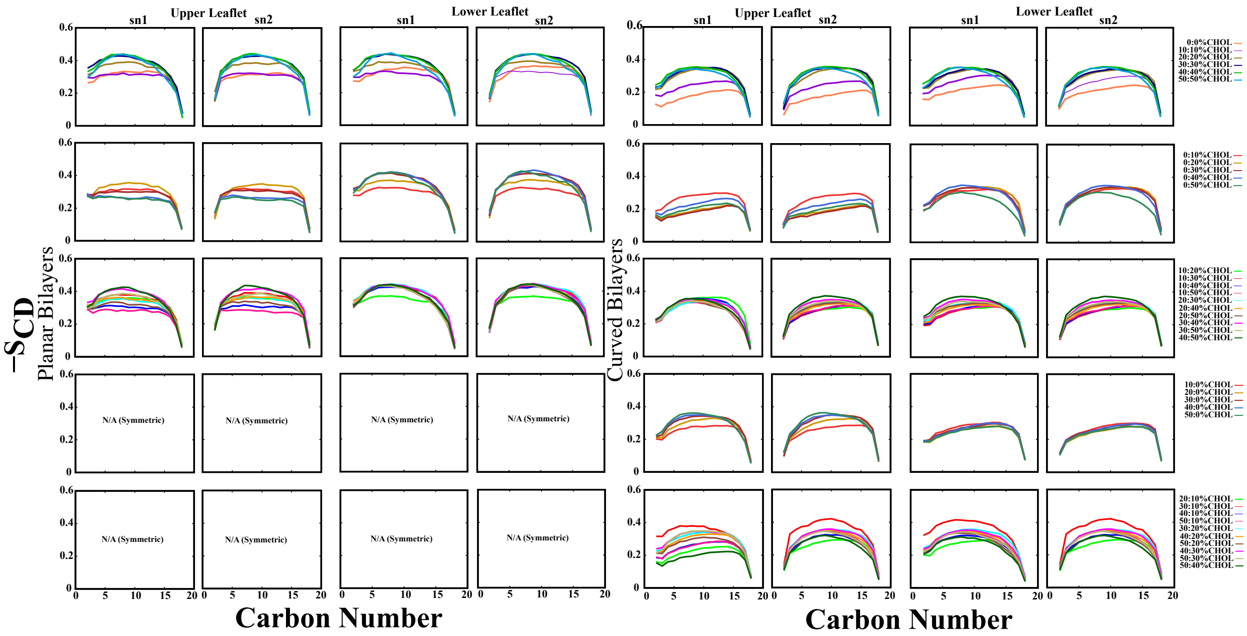

**Fig. S3. Segmental order parameter (*SCD*) profiles in planar and curved bilayers.** *SCD* profiles of sn1 and sn2 hydrocarbon chains in the upper and lower leaflets of both planar and curved bilayers with varying cholesterol concentrations. Each subplot corresponds to a specific leaflet and chain (sn1 or sn2), and each colored line represents a different cholesterol distribution (X–Y% CHOL), where X and Y denote the cholesterol percentages in the upper and lower leaflets, respectively. Higher *SCD* values indicate increased ordering and rigidity of the lipid tails. The data highlight how cholesterol concentration and asymmetry modulate lipid chain order in both geometries. Symmetric systems are marked as N/A where applicable.

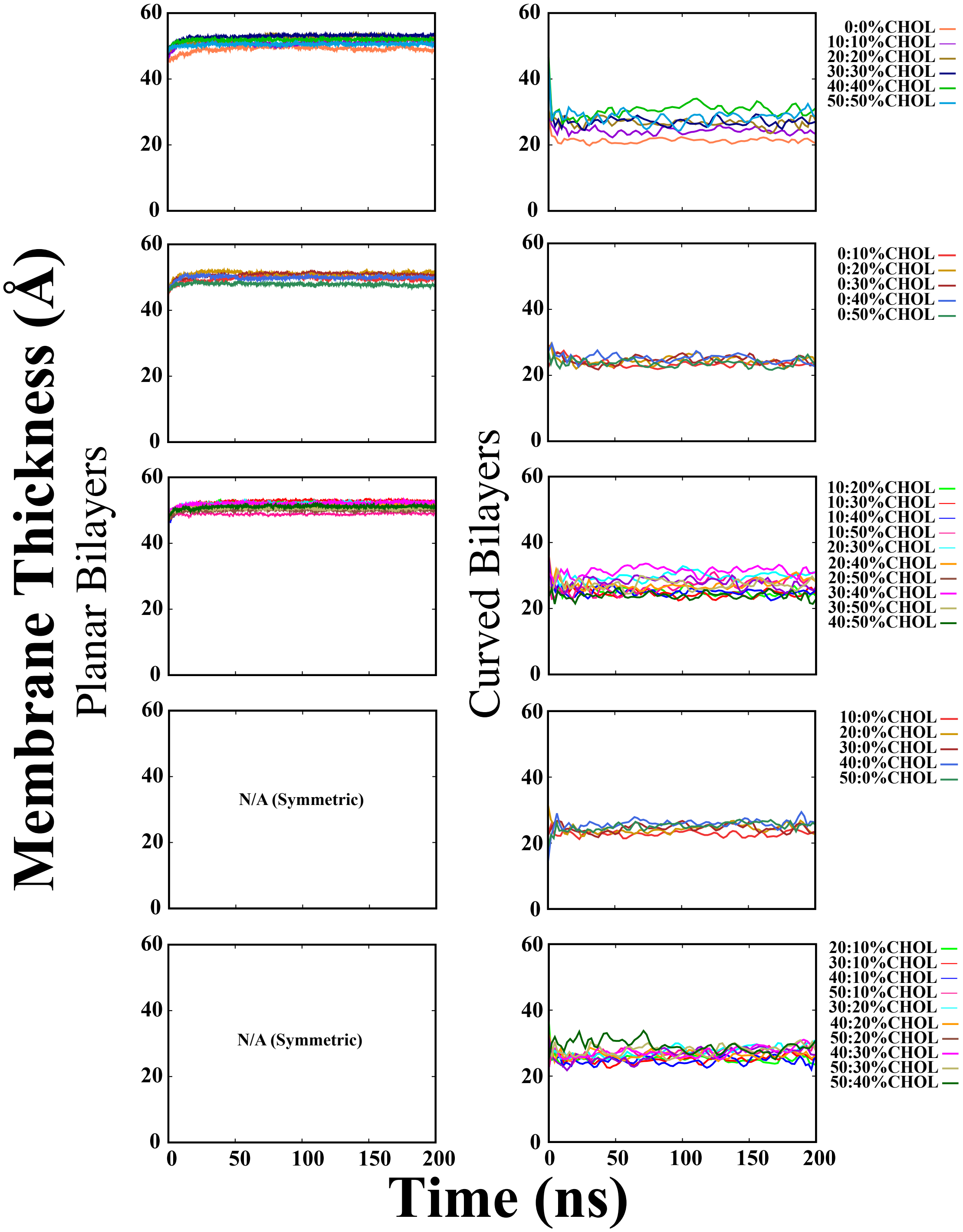


**Fig. S4. Membrane thickness in planar and curved bilayers.** Time series of the bilayer thickness over a 200 ns simulation under varying cholesterol conditions. Heatmaps show the average thickness across symmetric and asymmetric distributions, highlighting how curvature and cholesterol asymmetry modulate membrane structure.

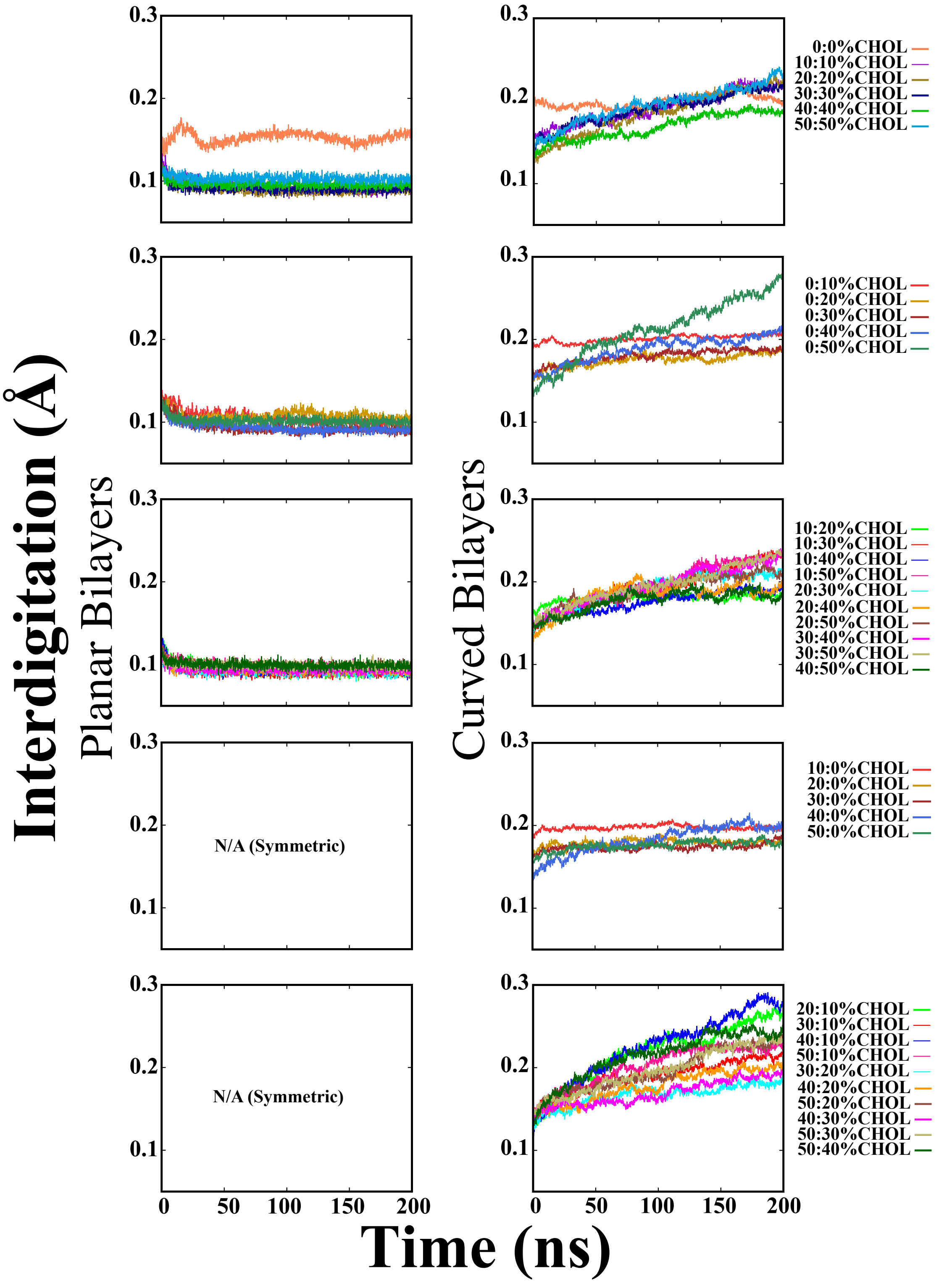


**Fig. S5. Lipid tail interdigitation in planar and curved DSPC:CHOL bilayers.** Time evolution of interdigitation (in Å) over 200 ns for systems with varying symmetric and asymmetric cholesterol concentrations. Each line represents a specific CHOL condition, labeled as X–Y% CHOL. Planar bilayers (left column) show low and stable interdigitation, while curved bilayers (right column) exhibit increasing interdigitation, especially in asymmetric and CHOL-rich systems. These results demonstrate how cholesterol distribution and curvature enhance interleaflet tail overlap.
